# Synaptic accumulation of GluN2B-containing NMDA receptors mediates the effects of BDNF-TrkB signalling on synaptic plasticity and in epileptogenesis

**DOI:** 10.1101/2024.10.21.618702

**Authors:** Pasqualino De Luca, Miranda Mele, Sara Tanqueiro, Francesca Napoli, Ugné Butkevičiūtė, Arthur C. Souto, Rui O. Costa, Alexander Schwarz, Meinrad Drexel, Ana M. Sebastião, Maria J. Diógenes, Carlos B. Duarte

**Affiliations:** CNC-Center for Neuroscience and Cell Biology, University of Coimbra, Portugal; Institute for Interdisciplinary Research, University of Coimbra, Coimbra, Portugal; PhD Programme in Experimental Biology and Biomedicine (PDBEB), Institute for Interdisciplinary Research (IIIUC), University of Coimbra, Portugal; Institute of Pharmacology and Neurosciences, Faculty of Medicine, and Unit of Neurosciences, Institute of Molecular Medicine, University of Lisbon, Lisbon, Portugal; Sechenov Institute of Evolutionary Physiology and Biochemistry of the Russian Academy of Sciences, St. Petersburg, Russia; Department of Pharmacology, Medical University of Innsbruck, Innsbruck, Austria; Department of Life Sciences, University of Coimbra, Coimbra, Portugal

## Abstract

Brain-derived neurotrophic factor (BDNF) is a key mediator of synaptic plasticity and memory formation in the hippocampus. However, the BDNF-induced alterations in the glutamate receptors coupled to the plasticity of glutamatergic synapses in the hippocampus have not been elucidated. In this work we investigated the putative role of GluN2B-containing NMDA receptors in the plasticity of glutamatergic synapses induced by BDNF. Stimulation of hippocampal synaptoneurosomes with BDNF led to a significant time-dependent increase in the synaptic surface expression of GluN2B-containing NMDA receptors as determined by immunocytochemistry with colocalization with pre- (vesicular glutamate transporter) and post-synaptic markers (PSD95). Similarly, BDNF induced the synaptic accumulation of GluN2B-containing NMDA receptors at the synapse in cultured hippocampal neurons, by a mechanism sensitive to the PKC inhibitor GӦ6983. The effects of PKC may be mediated by phosphorylation of Pyk2, as suggested by western blot experiments analyzing the phosphorylation of the kinase on Tyrosine 402. GluN2B-containing NMDA receptors mediated the effects of BDNF in the facilitation of the early phase of long-term potentiation (LTP) of hippocampal CA1 synapses induced by θ-burst stimulation, since the effect of the neurotrophin was abrogated in the presence of the GluN2B inhibitor Co 101244. In the absence of BDNF, the GluN2B inhibitor did not effect LTP. Surface accumulation of GluN2B-containing NMDA receptors was also observed in hippocampal synaptoneurosomes isolated from rats subjected to the pilocarpine model of temporal lobe epilepsy, after reaching Status epilepticus, an effect that was inhibited by administration of the TrkB receptor inhibitor ANA-12. Together, these results show that the synaptic accumulation of GluN2B-containing NMDA receptors mediate the effects of BDNF in the plasticity of glutamatergic synapses in the hippocampus.

## Introduction

Brain-derived neurotrophic factor (BDNF) is critical for long-term potentiation (LTP) induced by high-frequency presynaptic stimulation in the hippocampus, as well as in other brain regions (Minichiello 2009, Leal, Comprido et al. 2014, Costa, Martins et al. 2022). In addition, BDNF-TrkB signaling mediates the increase in neuronal excitability coupled to the development of seizures in experimental models of epilepsy (Gu, Huang et al. 2015). The activation of TrkB receptors by BDNF is coupled to the downstream activation of the Ras-ERK and PI3-K/Akt signaling pathways, as well as the phospholipase Cγ (Minichiello 2009, Leal, Comprido et al. 2014). In particular phospholipase Cγ is a key mediator of the effects of BDNF on hippocampal LTP, and a peptide uncoupling TrkB receptors from phospholipase Cγ1 prevented epilepsy induced by status epilepticus in mice (Gu, Huang et al. 2015). Local protein synthesis mediates the facilitatory effects of BDNF at CA1 hippocampal synapses (Kang and Schuman 1996, Schratt, Nigh et al. 2004, Takei, Inamura et al. 2004, Leal, Comprido et al. 2014, Costa, Martins et al. 2022), and is also required for consolidation of LTP upon infusion of the neurotrophin in the dentate gyrus (Messaoudi, Kanhema et al. 2007, Panja, Kenney et al. 2014). The postsynaptic proteins Arc (Yin, Edelman et al. 2002, Takei, Inamura et al. 2004), Homer2, Pyk2 and the Ca^2+^/calmodulin-dependent protein kinase II, together with the GluA1 AMPA receptor subunits are locally translated following synaptic stimulation with BDNF (Schratt, Nigh et al. 2004, Takei, Inamura et al. 2004, Afonso, De Luca et al. 2019). In addition to the effects of BDNF on local translation, the activity of the proteasome is also important in the early phase of LTP in CA1 synapses, providing a tight control of the synaptic proteome (Santos, Mele et al. 2015).

NMDA receptors (NMDAR) play important roles in the initiation of long-term synaptic plasticity, including long-term synaptic potentiation (LTP), which underlie certain forms of learning and memory. Alterations in NMDAR receptor activity have also been associated with different neurological and psychiatric disorders (Dupuis, Nicole et al. 2023). These receptors are formed by oligomerization of four subunits, and in most cases include two obligatory GluN1 subunits which bind glycine/D-serine and two GluN2 subunits that contain the glutamate binding site (Paoletti, Bellone et al. 2013, Wyllie, Livesey et al. 2013). The GluN2A and GluN2B subunits are the most abundant NMDAR subunits in the forebrain and in the CA1 region of the hippocampus, and the pattern of subunit composition determines the biophysical properties of the receptors, including the affinity for the agonist, open probability, Ca^2+^ permeability and charge transfer, and deactivation kinetics (Yashiro and Philpot 2008). These NMDAR subunits display a largely non-overlapping nanoscale distribution in hippocampal synapses, pointing to the activity of specific sorting mechanisms depending on the receptor composition (Zeng, Shang et al. 2016, Kellermayer, Ferreira et al. 2018).

The ratio between GluN2A and GluN2B subunits is different between synapses and may change depending on the development stage, in response to synaptic stimulation and in association with axonal refinement following sensory experience (Yashiro and Philpot 2008, Carta, Srikumar et al. 2018, Dupuis, Nicole et al. 2023). Together, these findings point to the availability of regulatory mechanisms that change the composition of GluN2-containing NMDAR to modulate their function based on the physiological context. For example, an impairment in LTP of commissural-CA3 synapses was observed in experiments performed in conditional knock-out mice in which GluN2B subunits were downregulated specifically in hippocampal CA3 pyramidal neurons (Akashi, Kakizaki et al. 2009). On the other hand, overexpression of GluN2B in the forebrain of transgenic mice facilitated the potentiation of CA1 synapses in response to high frequency stimulation, and the effects were associated with improved learning and memory in different tasks (Tang, Shimizu et al. 1999). Studies performed using the pilocarpine model of temporal lobe epilepsy, when the animals display spontaneous seizures, also showed an upregulation in GluN2B-mediated responses in the lateral perforant path synapses with hippocampal dentate granule neurons (Klatte, Kirschstein et al. 2013), and similar changes account for the facilitation of NMDAR-dependent LTP of Schaffer collateral-CA1 synapses (Muller, Tokay et al. 2013, Gorlewicz, Pijet et al. 2022).

The surface NMDAR travel between the synaptic and extrasynaptic sites, with more than one third of the receptors displaying an extrasynaptic location (Dupuis, Nicole et al. 2023). The available evidence shows that GluN2B-containing NMDAR are more mobile than GluN2A, which are preferentially located at the synapse (Groc, Heine et al. 2006, Dupuis, Ladepeche et al. 2014, Ferreira, Papouin et al. 2017, Kellermayer, Ferreira et al. 2018). The pool of surface NMDAR is dictated by the rate of synthesis in the endoplasmic reticulum, their dendritic transport, exocytosis to the cell surface, lateral diffusion within the plasma membrane, and the rates of endocytosis, recycling and degradation. The C-terminus varies significantly between NMDAR subunits and play a key role in the regulation of the receptor channel properties, trafficking and in the activation of downstream signaling. These functions can be differentially regulated by alternative splicing, interaction with binding partners and post-translational modifications (Warnet, Bakke Krog et al. 2021). Local protein synthesis was also shown to control indirectly the synaptic surface expression of GluN2B-containing NMDAR (Leal, Comprido et al. 2017, Afonso, De Luca et al. 2019).

Although different forms of synaptic plasticity have been shown to be associated with the synaptic recruitment of GluN2B-containing NMDAR, the underlying molecular mechanisms have not been elucidated. In this work we have elucidated a key role for BDNF-TrkB signaling in the upregulation of GluN2B function and localization in hippocampal synapses associated with the facilitation of LTP and with hyperexcitability during status epilepticus, an action that involves synaptic accumulation of GluN2B mediated by activation of protein kinase C.

## Material and Methods

### Animals

Wistar rats and male Sprague-Dawley rats, aged 6-8 weeks (Charles River Laboratories, Barcelona, Spain), were maintained under controlled temperature (21 ± 1°C) and humidity (55 ± 10%) conditions, with a 12:12 h light/dark cycle and access to food and water ad libitum. All animals were handled according to Portuguese Law (DL 113/2013) and the European Community Guidelines (Directive, 2010/63/EU) on the protection of animals for scientific experimentation.

### Isolation of synaptoneurosomes

Synaptoneurosomes were isolated as previously described with slight modifications (Afonso, De Luca et al. 2019, Masella, Silva et al. 2024). Briefly, 2 hippocampi were dissected from adult Sprague-Dawley rats (6-8 weeks old). The tissue was minced with a blade and homogenized with Kontes Dounce Tissue Grinder in a buffer containing 0.32 M sucrose, 10 mM HEPES-Tris pH 7.4, and 0.1 mM EGTA pH 8, using first a pestle 0.889-0.165 mm clearance, followed by a pestle of 0.025-0.076 mm. After centrifugation for 3 min at 1000 x g at 4°C, the collected supernatant was passed through triple layered nylon membranes (150 and 50 μm, VWR) and then through an 8 μm pore size filter (Millipore, MA). The flow-through was centrifuged for 15 min at 10000 x g, and the resulting pellet was suspended in the buffer used for the homogenization. All the procedure was performed on ice or at 4 °C. Part of the synaptoneurosome preparation was plated on glass coverslips and processed for immunostaining, the remaining material was used for protein extraction followed by western blot analyses (see below).

### Hippocampal cultures

Neuronal low-density hippocampal cultures were prepared from the hippocampi of E18-E19 Wistar rat embryos, after incubation with trypsin (0.06% (w/v), GIBCO - Life Technologies) in Ca^2+^- and Mg^2+^-free Hanks’ balanced salt solution (HBSS: 5.36 mМ KCl, 0.44 mM KH_2_PO_4_, 137 mM NaCl, 4.16 mM NaHCO_3_, 0.34 mM Na_2_HPO_4_·2H_2_O, 5 mM glucose, 1 mM sodium pyruvate, 10 mM HEPES, and 0.001% phenol red), for 15 min at 37°C (Afonso, De Luca et al. 2019). The hippocampi were then washed with Hanks’ balanced salt solution containing 10% fetal bovine serum (GIBCO - Life Technologies) to stop trypsin activity and transferred to Neurobasal medium (GIBCO - Life Technologies) supplemented with SM1 supplement (1:50 dilution, STEMCELL Technologies), 25 μM glutamate, 0.5 mM glutamine, and 0.12 mg/ml gentamycin (GIBCO - Life Technologies). Neurons were plated at a final density of 1.4 x 10^4^ cells/cm^2^ on poly-D-lysine-coated glass coverslips in neuronal plating medium (Minimum Essential Medium [MEM, Sigma-Aldrich] supplemented with 10% horse serum, 0.6% glucose and 1 mM pyruvic acid). After 2-4 h the coverslips were flipped over an astroglia feeder layer in Neurobasal medium (GIBCO - Life Technologies) supplemented with SM1 supplement (1:50 dilution, STEMCELL Technologies), 25 μM glutamate, 0.5 mM glutamine and 0.12 mg/ml gentamycin (GIBCO - Life Technologies). The neurons grew face down over the feeder layer but were kept separate from the glia by wax dots on the neuronal side of the coverslips. To prevent the overgrowth of glial cells, neuronal cultures were treated with 10 μM 5-Fluoro-2′-deoxyuridine (Sigma-Aldrich) at DIV 3. Cultures were maintained in a humidified incubator with 5% CO_2_/95% air at 37°C for up to 2 weeks, feeding the cells once per week with the same Neurobasal medium described above, but without glutamate added. At DIV 14-15 the neurons were stimulated with 50 ng/ml BDNF (Peprotech) for the indicated period of time. Where indicated, cells were pre-treated for 45 min with a protein synthesis inhibitor, 40-45 min with a protein kinase C inhibitor, (3-[1-[3-(Dimethylamino)-propyl]-5-methoxy-1H-indol-3-yl]-4-(1H-indol-3-yl)-1H-pyrrole-2,5-dione (GӦ6983, 100 nM) or with the vehicle dimethyl sulfoxide (DMSO, 1:1000 dilution, Sigma-Aldrich), as control. High-density hippocampal cultures were prepared from the hippocampi of E18-E19 Wistar rat embryos, as described above after treatment with trypsin as described above. After dissociation in Neurobasal medium (Gibco-Life Technologies) supplemented with SM1 supplement (1:50 dilution; STEMCELL Technologies), 25 µM glutamate, 0.5 mM glutamine, and gentamicin (0.12 mg/ml) (Gibco-Life Technologies), the cells were plated in six-well plates (85.5×10^3^ cells/cm^2^) coated with poly-D-lysine (0.1 mg/ml) for biochemical purposes (Western blot). Cultures were maintained in a humidified incubator with an atmosphere of 5% CO_2_/95% air at 37°C for 14 days, and were then stimulated with BDNF (50 ng/ml) (PeproTech) for the indicated periods.

### Preparation of hippocampal slices

For electrophysiological studies, male Wistar rats (7–8 weeks old) were sacrificed after being deeply anesthetized with isoflurane (Esteve). The brain was quickly removed and placed in ice-cold oxygenated (95% O_2_/5% CO_2_) artificial cerebrospinal fluid (aCSF) (124 mM NaCl, 3 mM KCl, 1.2 mM NaH_2_PO_4_, 25 mM NaHCO_3_, 2 mM CaCl_2_, 1 mM MgSO_4_, and 10 mM glucose, pH 7.4) and both hippocampi were dissected. Hippocampal slices were cut perpendicularly to the long axis of the hippocampus (400 μm thick) with a McIlwain tissue chopper. After recovering functionally and energetically for at least 1 h in a resting chamber filled with oxygenated aCSF at RT, hippocampal slices were transferred to a recording chamber continuously superfused with oxygenated aCSF at 32°C (flow rate of 3 mL/min).

### Staining of surface GluN2B-containing NMDAR in cultured hippocampal neurons and quantitative image analysis

To label surface GluN2B-containing NMDAR, live neurons (low-density hippocampal cultures) were incubated for 10 min at room temperature (RT) with an antibody against an extracellular epitope of the GluN2B N-terminus (1:100; AGC-003, Alomone Labs) diluted in a saline buffer (145 mM NaCl, 5 mM glucose, 10 mM HEPES, 5 mM KCl, 1.8 mM CaCl_2_, 1 mM MgCl_2,_ [pH 7.3]) (Sham) solution, as previously described (Afonso, De Luca et al. 2019). Neurons were then fixed for 15 min in 4% sucrose and 4% paraformaldehyde in phosphate-buffered saline (PBS: 137 mM NaCl, 2.7 mM KCl, 1.8 mM KH2PO_4_, 10 mM Na_2_HPO_4_·2H_2_O [pH 7.4]) at RT and permeabilized with PBS + 0.3% (v/v) Triton X-100 for 5 min at 4°C. The preparations were then incubated in 10% (w/v) bovine serum albumin (BSA) in PBS for 1 h at RT to block nonspecific staining and incubated with the appropriate primary antibody (anti-MAP2 [1:10,000; ab5392, Abcam], anti–PSD-95 [1:200; 7E3-1B8, Thermo Scientific], and anti-vGluT1 [1:5000; AB5905, Millipore]) diluted in 3% (w/v) BSA in PBS (overnight, 4°C). After washing six to seven times in PBS, cells were incubated with the appropriate secondary antibody (Alexa Fluor 488–conjugated anti-mouse [1:1000; A-11001, Thermo Fisher Scientific], Alexa Fluor 488-conjugated anti-rabbit [1:1000; A-11034, Thermo Fisher Scientific], Alexa Fluor 568–conjugated anti-mouse [1:1000; A-11004, Thermo Fisher Scientific], Alexa Fluor 568-conjugated anti-rabbit [1:1000; A-11036, Thermo Fisher Scientific], Alexa Fluor 647-conjugated anti– guinea pig [1:500; A-21450, Thermo Fisher Scientific], and AMCA (aminomethyl-coumarin)-conjugated anti-chicken [1:200; #103-155-155, Jackson ImmunoResearch]) diluted in 3% (w/v) BSA in PBS (1 h at RT). The coverslips were mounted using a fluorescent mounting medium (DAKO).

Fluorescence imaging was performed on a Zeiss Axio Observer Z.1 microscope using a 63 × 1.4 numerical aperture (NA) oil objective and a Zeiss Axio Imager Z.2 microscope using a 63× 1.4 NA oil objective, both equipped with a Zeiss HRm AxioCam. Images were quantified using the Fiji image analysis software. For quantification, sets of cells were cultured and stained simultaneously and imaged using identical settings. The region of interest was randomly selected, and the dendritic length was measured based on MAP2 staining. The protein signals were analyzed after setting the appropriate thresholds, and the recognizable puncta under those conditions were included in the analysis. For each experiment, similar threshold levels were used to quantify the number and the integrated intensity of puncta in dendrites. Measurements were performed in three to six independent preparations, as indicated in the figure captions.

To analyze GluN2B synaptic surface expression, the PSD-95 and vGluT1 signals were thresholded and their colocalization was determined. The surface GluN2B signal was measured after thresholds were set so that recognizable puncta were included in the analysis. The surface GluN2B signal present in glutamatergic synapses was obtained by measuring the surface GluN2B puncta positive for both PSD-95 and vGluT1. The results were represented *per* density of excitatory synapses (number of positive PSD-95–vGlut1 puncta that colocalized per dendritic length). To quantify the synaptic surface GluN2B immunoreactivity, digital images were subjected to a user-defined intensity threshold, to select puncta and measured for puncta intensity and number, for the selected region. The synaptic GluN2B puncta were identified by their overlap with the thresholded PSD-95 signal. The results were represented per dendritic length.

### Live immunostaining of surface GluN2B-containing NMDAR in hippocampal synaptoneurosomes

Synaptoneurosomes (35 µl) were plated on 10 mm diameter coverslips coated with poly-D-lysine (0.1 mg/mL) and left in a humid chamber at RT for 1 h, to allow adhesion. Where indicated, synaptoneurosomes were incubated with BDNF (50 ng/mL, Peprotech) at 30⁰C during this period, and control experiments in the absence of the neurotrophin were also performed. Immediately after stimulation, synaptoneurosomes were processed for GluN2B live immunostaining as described below (see also (Masella, Silva et al. 2024)). Live staining was performed as described above for the analysis of surface GluN2B expression in cultured hippocampal neurons (Afonso, De Luca et al. 2019, Masella, Silva et al. 2024).

### Fluorescence imaging acquisition and quantitative analysis of NMDAR surface staining in synaptoneurosomes

Fluorescence imaging of synaptoneurosomes was conducted using a Carl Zeiss Axio Imager Z2 widefield fluorescence microscope, following the protocol described in (Masella, Silva et al. 2024). Images were captured with a Plan-Apochromat 100×/1.4 numerical aperture oil objective and a Zeiss HRm AxioCam. The imaging utilized Zeiss filter sets 31, 38 (HE), and 50, and phase-contrast images were also obtained. Consistent exposure times and light intensities were maintained across different experimental conditions. Image analysis was performed with ImageJ software (National Institutes of Health, MD, USA) using a custom-designed macro for this experiment. Intact synaptoneurosomes were identified based on the following criteria: juxtaposition of presynaptic (VGluT1) and postsynaptic (PSD-95) protein clusters, visibility as dark objects in phase-contrast images indicating sealed synaptoneurosomes, size range (300–1000 nm), and a snowman-like shape as detailed in reference (Tai, Wang et al. 2014). Approximately 500–600 synaptoneurosomes per condition were manually delineated as regions of interest (ROIs), and background and thresholds were set for GluN2B signals. The analysis quantified the area mean intensity as well as the integrated density of GluN2B, staining within the synaptoneurosomes.

### Preparation of extracts from cultured hippocampal neurons and hippocampal synaptoneurosomes

Hippocampal cultures at 15 DIV (85.5 × 10^3^ cells/cm^2^) were washed twice with ice-cold PBS and once more with PBS buffer supplemented with 1 mM DTT and a cocktail of protease inhibitors (0.1 mM phenylmethylsulfonyl fluoride [PMSF] and CLAP [chymostatin (1 µg/ml), leupeptin (1 µg/ml), antipain (1 µg/ml), and pepstatin (1 µg/ml); Sigma]). The cells were then lysed with RIPA buffer (150 mM NaCl, 50 mM Tris-HCl, 5 mM EGTA, 1 % Triton X-100, 0.5 % deoxycholic acid, 0.1 % sodium dodecyl sulfate) supplemented with 1.5 mM sodium orthovanadate (Na_3_VO_4_), 50 mM NaF and the cocktail of protease inhibitors. The RIPA buffer was also supplemented with 1.5 mM sodium orthovanadate in the preparation of synaptoneurosomal extracts. The extracts were frozen at −80°C, defrosted, and centrifuged at 16,000 × g for 10 min. Protein in the supernatants was quantified using the BCA assay kit (Pierce). In the analysis of extracts from cultured hippocampal neurons, the samples were denatured with 2x concentrated denaturing buffer (125 mM Tris [pH 6.8], 100 mM glycine, 4% SDS, 200 mM DTT, 40% glycerol, 3 mM Na_3_VO_4_, and 0.01% bromophenol blue) for 5 min at 95°C, and proteins of interest were analyzed by Western blot. Synaptoneurosomal samples were diluted with Laemmli Sample Buffer (Bio-Rad) supplemented with 5 % β-mercaptoethanol.

### Western blotting

Proteins were separated by SDS-PAGE in 10% polyacrylamide gels and transferred to polyvinylidene difluoride membrane (Millipore) by electroblotting (40 V, overnight at 4°C). In the experiments analyzing Pyk2 and pPyk2 protein levels, the membranes were blocked for 1 h with 5% skim milk in TBS-T (20 mM Tris, 137 mM NaCl [pH 7.6], supplemented with 0.1% Tween 20), and probed with the primary antibody overnight at 4°C (anti-Pyk2 [1:500; sc-1514, Santa Cruz Biotechnology] and anti-pPyk2 [1:500; 44-618G, Invitrogen]). After several washes with TBS-T, the membranes were incubated with an alkaline phosphatase-conjugated immunoglobulin G (IgG) secondary antibody (anti-rabbit or anti-goat, depending on the primary antibody host-species; Jackson ImmunoResearch) for 1 h at RT, washed again and incubated with enhanced chemifluorescent substrate, ECF (GE Heathcare), for up to 5 min at RT. For detection of TrkB and pTrkB in hippocampal synaptoneurosomes, the membranes were blocked in 3 % BSA in TBS-T (20 nM Tris, 13.7 mM NaCl, 0.1 % Tween, pH 7.6) before incubation with the primary antibodies diluted in 1 % BSA TBS-T (overnight at 4°C). The following primary antibodies were used: anti-phospho-TrkB (Tyr816) (1:1000; Sigma-Aldrich) and anti-TrkB (1:1000; BD Transduction Laboratories). The anti-β-tubulin (1:200.000, Sigma) antibody was used as loading control. Finally, the membranes were washed and exposed to StarBright Blue 520 and 700 fluorophores-conjugated secondary antibodies (Goat Anti-rabbit IgG StarBright Blue 700 and Goat Anti-mouse IgG StarBright Blue 520 (Bio-Rad), 1:5,000 dilution; 1 h at RT). Immunostaining was visualized by the enhanced chemifluorescence method on ChemiDoc (BioRad). Quantification of the signals was performed with the Fiji image analysis software.

### *Ex Vivo* electrophysiology recordings in hippocampal slices

Field excitatory postsynaptic potentials (fEPSPs) were recorded extracellularly through a microelectrode filled with aCSF (2–6 MΩ) placed in the stratum radiatum of the CA1 area. The stimulation (rectangular 0.1 ms pulses, once every 10 s) was delivered alternatively to two independent pathways through bipolar concentric wire electrodes placed on the Schaffer collateral fibers as previously described (Diogenes, Costenla et al. 2011). Recordings were obtained with an Axoclamp 2B amplifier (Axon Instruments, Foster City, CA, United States), digitized and continuously stored on a personal computer with the WinLTP 2.20b software (Andersson, Stenqvist et al. 2001). Individual responses were monitored and averages of six consecutive responses were obtained and the slope of the initial phase of the fEPSP was quantified. LTP was induced by a θ-burst protocol (four trains of 100 Hz, four stimuli, separated by 200 ms) since the facilitatory action of BDNF upon LTP is mostly seen under this paradigm (Diogenes, Costenla et al. 2011). In the control experiments, LTP was inducted in the first pathway after obtaining a stable fEPSP slope. After 60 min of LTP induction, BDNF (20 ng/mL) was added to the superfusion solution. Whenever necessary, the intensity of stimulation was adjusted to obtain fEPSP slopes of similar values as recorded before BDNF application. After at least 20 min of BDNF perfusion, and a stable baseline of fEPSP slope values was obtained, LTP was induced in the second pathway. Each pathway was used as control (to be used to induce the first LTP) or test (for the second LTP) on alternate days. When testing the effect of Co 101244 (Tocris) upon LTP, Co 101244 (1 µM) was added to the slices at least 20 min before the second LTP induction. To test the modulatory effect of Co 101244 over the effect of BDNF upon LTP, that drug was added to the superfusion solution at least 20 min before LTP induction in the first pathway and remained in the bath till the end of the experiment. LTP was quantified as a percentage of change in the average slope of the fEPSP taken from 46 to 60 min after LTP induction relatively to the average slope of the fEPSP measured during the 10 min before LTP induction.

### Lithium-pilocarpine model of Status Epilepticus

The protocol used to induce status epilepticus using the lithium-pilocarpine model was approved by DGAV-Direcção Geral de Ambiente e Veterinária (Ref. 0421/000/000/2020). Experiments were conducted using 6-8 weeks-old male Sprague-Dawley rats obtained from Charles River Laboratories. First, the animals were weighed to determine the correct dosage of each substance. Lithium chloride (LiCl; 127 mg/ml in deionized sterile water) was administered first at a dosage of 127 mg/kg via intraperitoneal injection. Twenty hours later, methyl-scopolamine (2 mg/mL in saline [0.9% NaCl]) was administered at a dosage of 1 mg/kg, also via intraperitoneal injection. The animals were maintained for 45-min more before intraperitoneal injection of pilocarpine (20 mg/mL in saline [0.9% NaCl]) at a dosage of 10 mg/kg. During this period the animals were closely monitored for any adverse reactions. Behavioural changes and physiological responses were recorded with a video camera, with the severity of seizures classified using the Racine scale (Racine 1972) that categorizes the progression of seizure severity based on observable behaviours. If an animal did not reach Racine stage 5, indicative of status epilepticus (SE), within 30 min of the pilocarpine injection, an additional dose of pilocarpine (10 mg/kg) was administered. This procedure was repeated every 30 min, up to a maximum of five times, until SE is achieved. A saline control group followed the same protocol as the pilocarpine-injected group, but with saline injections replacing pilocarpine administration at equivalent volumes.

For experiments performed in the presence of the compound ANA-12, the drug was injected (i.p.) at a concentration of 0.5 mg/mL (prepared in saline, 5% DMSO) one hour before the first pilocarpine injection. ANA-12 was administered at a dosage of 0.5 mg/kg. Following this procedure, the animals were maintained for 15-min before administration of methyl-scopolamine, and pilocarpine was administered 45 min later. A vehicle control group received an equivalent volume of a saline/DMSO solution, administered intraperitoneally, and based on the volume used for injection of ANA-12. Rats were anesthetized with isoflurane and euthanized by decapitation, 90 min after the onset of SE determined by the rat’s entry into the Stage 5 of seizures according to the Racine scale. The hippocampi were then dissected and processed for synaptoneurosome isolation as described above.

### Statistical analysis

The results are presented as means ± SEM. Statistical analysis was performed using parametric tests. Student’s *t-test* was used to compare two groups of samples. Comparisons between multiple groups were performed as indicated in the figure captions, using the Prism 8 software (GraphPad).

## Results

### BDNF increases the expression of GluN2B-containing NMDAR in hippocampal synaptoneurosomes

BDNF upregulates the activity of postsynaptic NMDAR in cortical and hippocampal CA1 pyramidal neurons, in dentate gyrus granule cells (Kolb, Trettel et al. 2005, Madara and Levine 2008), as well as in cultured hippocampal neurons (Leal, Comprido et al. 2017, Afonso, De Luca et al. 2019). The studies performed in cultured hippocampal neurons showed that the upregulation in NMDAR activity is associated with an enhanced synaptic accumulation of the receptors (Leal, Comprido et al. 2017, Afonso, De Luca et al. 2019), but it remains to be determined whether BDNF has the same effect in synapses from adult animals. This question was addressed using hippocampal synaptoneurosomes, a subcellular fraction containing the pre- and postsynaptic components, isolated from adult rats. Synaptoneurosomes were stimulated with 50 ng/mL BDNF for 10, 30, and 60 min before live immunostaining with an antibody against the extracellular N-terminus of GluB2B, which allows labeling surface receptors (Fig. 1A). Synaptic surface GluN2B-containing NMDAR were identified by colocalization with the pre-and post-synaptic markers vesicular glutamate transporter 1 (vGluT1) and PSD-95, respectively. The results were analyzed for the integrated density (Fig. 1B) and mean gray value (Fig. 1C) of GluN2B immunoreactivity that colocalized with the synaptic markers. BDNF up-regulated the integrated density (Fig. 1B) and mean gray value (Fig. 1C) of GluN2B immunoreactivity in hippocampal synaptoneurosomes stimulated for 10, 30, and 60 min. Together, these results showed that BDNF induced a specific increase of surface GluN2B-containing NMDAR in the hippocampal synapses from adult rats.

**Figure 1.**
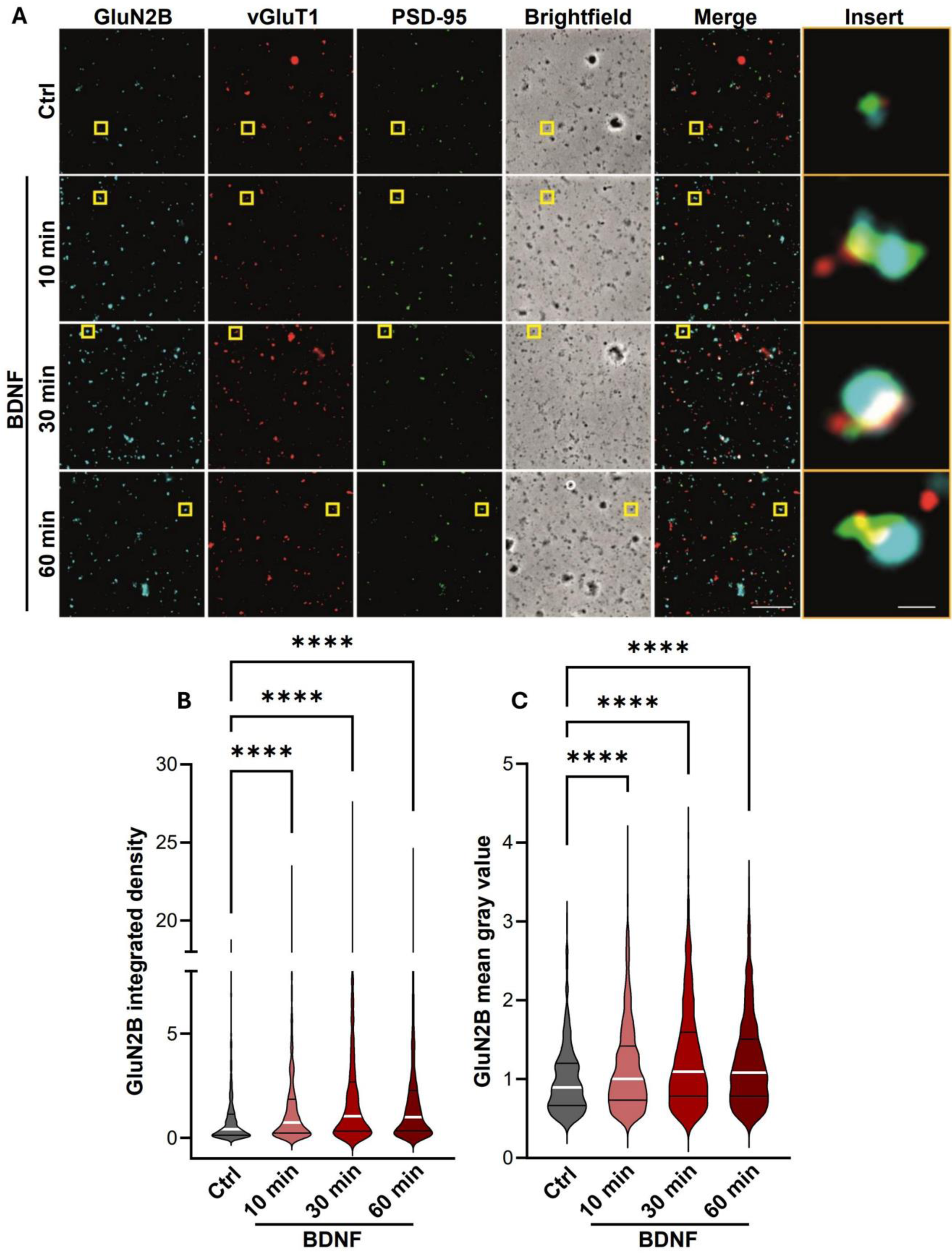
BDNF-induced increase in the expression of GluN2B-containing NMDAR in hippocampal synaptoneurosomes. (**A**) Representative images of hippocampal synaptoneurosomes (prepared from adult Sprague-Dawley rats 6-8 weeks old) incubated with BDNF (50 ng/mL). Synaptoneurosomes were immunoassayed for GluN2B using an antibody against an extracellular epitope in the GluN2B N-terminus and immunoassayed for vGlut1, PSD-95. Merge scale bar, 10 μm. Insert scale bar, 0.5 µm. Images illustrated in (**A**) were analyzed for the GluN2B integrated density (**B**) and GluN2B mean gray value (**C**). Data are the means ± SEM of 779 - 840 synaptoneurosomes per condition, in at least three independent experiments performed in different preparations. ****p < 0.0001, by Kruskal-Wallis’s test and Dunn’s multiple comparisons test.

We then compared the time-dependency of synaptic surface accumulation of GluN2B in rat hippocampal synaptoneurosomes (Fig. 1) and synaptic regions of cultured hippocampal neurons (Fig. 2). In the latter preparation, surface GluN2B-containing NMDAR were live-stained with an antibody against an extracellular epitope of GluN2B and after fixation and permeabilization of the plasma membrane the cells were incubated with antibodies against the dendritic marker microtubule-associated protein 2 (MAP2), and the pre- and post-synaptic markers vGluT1 and PSD-95, respectively. Cultured hippocampal neurons were analyzed for the number (Fig. 2A, B), area (Fig. 2A, C) and intensity (Fig. 2A, D) of GluN2B puncta per dendritic length (as determined based on MAP2 staining). The effect of BDNF on the synaptic surface abundance of GluN2B was assessed by quantifying the number (Fig. 2A, E), area (Fig. 2A, F) and intensity (Fig. 2A, G) of puncta that co-localized with PSD-95 and vGluT1. Neurons incubated with BDNF for 10 min showed no significant changes in the total and synaptic surface abundance of the GluN2B subunit in dendrites (Fig. 2B-2G), in contrast with what was observed in hippocampal synaptoneurosomes (Fig. 1). However, incubation of hippocampal neurons with BDNF for 30 min increased the total GluN2B number of dendritic puncta (Fig. 2B), as well as in their area (Fig. 2C), and similar results were obtained for the surface synaptic GluN2B staining. Analysis of the synaptic surface GluN2B, also showed a significant increase in the intensity of staining after incubation of hippocampal neurons with BDNF for 30 min (Fig. 2G). These results showed that BDNF increases the synaptic surface expression of GluN2B-containing NMDARs in cultured hippocampal neurons and in rat hippocampal synaptoneurosomes, the action being even faster in synaptoneurosomes isolated from mature neurons than in cultured neurons.

**Figure 2.**
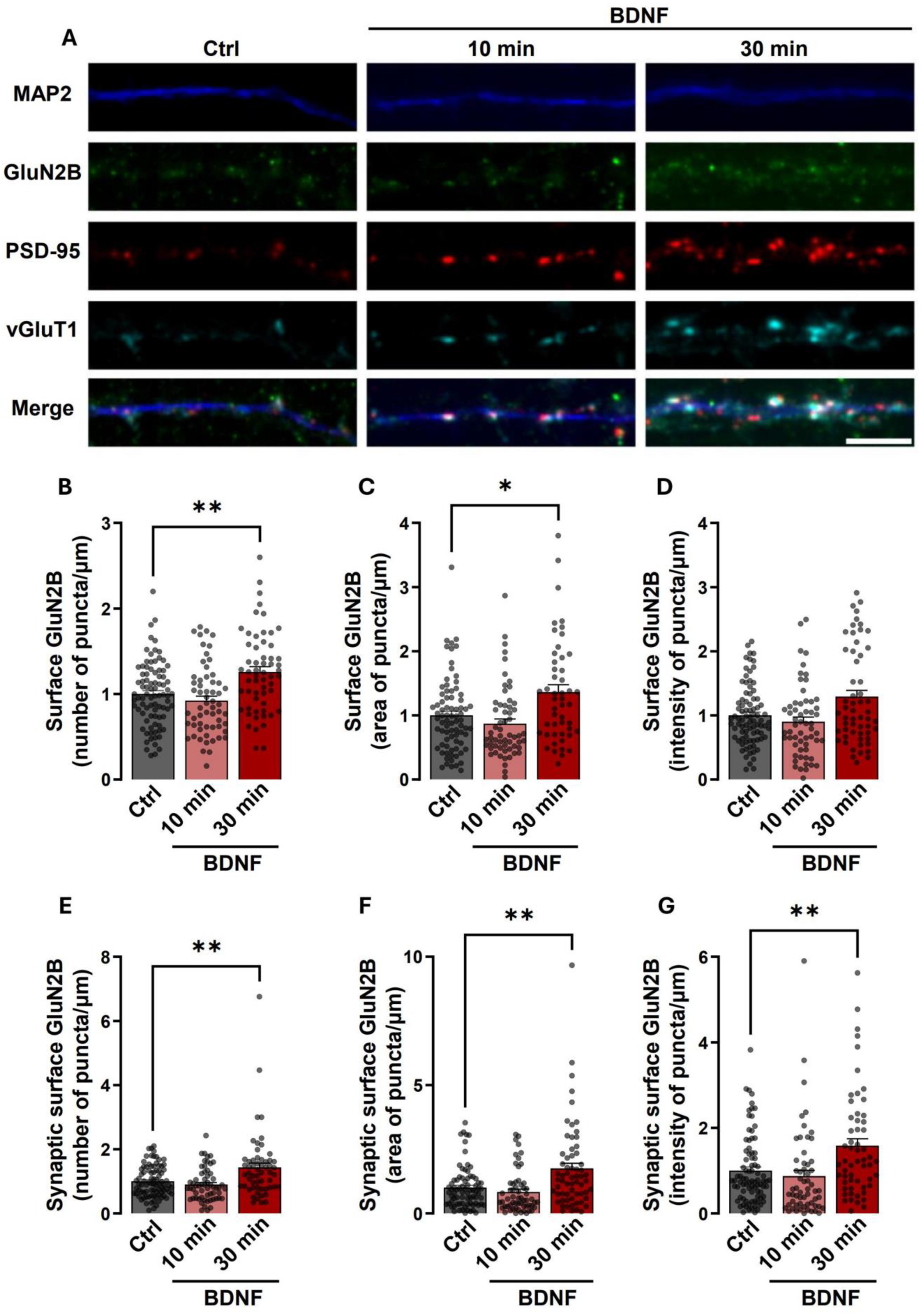
Stimulation of hippocampal neurons with BDNF for 30 min increases the synaptic expression of GluN2B-containing NMDAR. (**A**) Representative images of hippocampal neurons (DIV 14 - 15) that were stimulated with BDNF (50 ng/ml for 10 or 30 min), as indicated. Neurons were then live-immunoassayed for GluN2B using an antibody against an extracellular epitope in the GluN2B N-terminus, fixed, and then further immunoassayed for PSD-95, vGlut1, and MAP2. Scale bar, 5 μm. Images illustrated in (**A)**, were analyzed for the total number (**B**), intensity (**C**), and area (**D**) of surface GluN2B puncta per µm. Synaptic (PSD-95- and vGlut1-colocalized) surface GluN2B number (**E**), area (**F**), and intensity (**G**) of puncta per density of excitatory synapses (number of puncta PSD-95–vGlut1 colocalized per µm), were also analyzed. Data are normalized to the means of the control and are the means ± SEM of 43-45 cells per condition, from at least three independent experiments performed in different preparations. *p < 0.05, **p < 0.01 by one-way analysis of variance (ANOVA) followed by Bonferroni post-test.

### BDNF increases synaptic expression of GluN2B-containing NMDAR through a PKC-dependent mechanism

Local synthesis of the non-receptor tyrosine kinase Pyk2 was shown to mediate the effects of BDNF on the synaptic accumulation of GluN2B-containing NMDAR in cultured hippocampal neurons (Afonso, De Luca et al. 2019), but the signaling mechanisms that act downstream of TrkB receptors to activate Pyk2 remain to be elucidated. Previous studies performed in different preparations showed that Pyk2 activation can be triggered by stimulating PKC (Das, Pal et al. 2021, de Pins, Mendes et al. 2021), a kinase that is stimulated after TrkB-mediated activation of phospholipase C-γ (Leal, Comprido et al. 2014). Therefore, we hypothesized that this signalling pathway could mediate the activation of Pyk2 following stimulation of hippocampal neurons with BDNF, with a concomitant upregulation of GluN2B synaptic surface expression. To investigate the effect of BNDF in the activation of Pyk2, we analysed, by western blot, the phosphorylation of the kinase on Tyr402 in cultured hippocampal neurons. Phosphorylation of this residue plays an important role in the activation of downstream signalling (Das, Pal et al. 2021, de Pins, Mendes et al. 2021). BDNF (50 ng/ml) significantly increased total levels of phosphorylated Pyk2 (pPyk2) after 30 min of stimulation compared to the control condition (Fig 3A, 3C), with no effect on the total abundance of the protein (Fig. 3A, 3D). The membranes were then reprobed with an antibody against total Pyk2 (Fig 3A, 3D), and the mean densities of pPyk2(pY402) were expressed as ratios of the corresponding Pyk2 (total) (Fig 3A, 3B). Stimulation of hippocampal neurons with BDNF for 30 min also increased the ratio pPyk2(pY402)/Pyk2, while no effects were observed in experiments testing the effects of the neurotrophin for 10 min (Fig 3A - 3D). These results demonstrated that BDNF upregulates pPyk2(pY402) protein levels within a time frame that is compatible with a role for the kinase in the recruitment of GluN2B-containing NMDAR to the synaptic surface.

**Figure 3.**
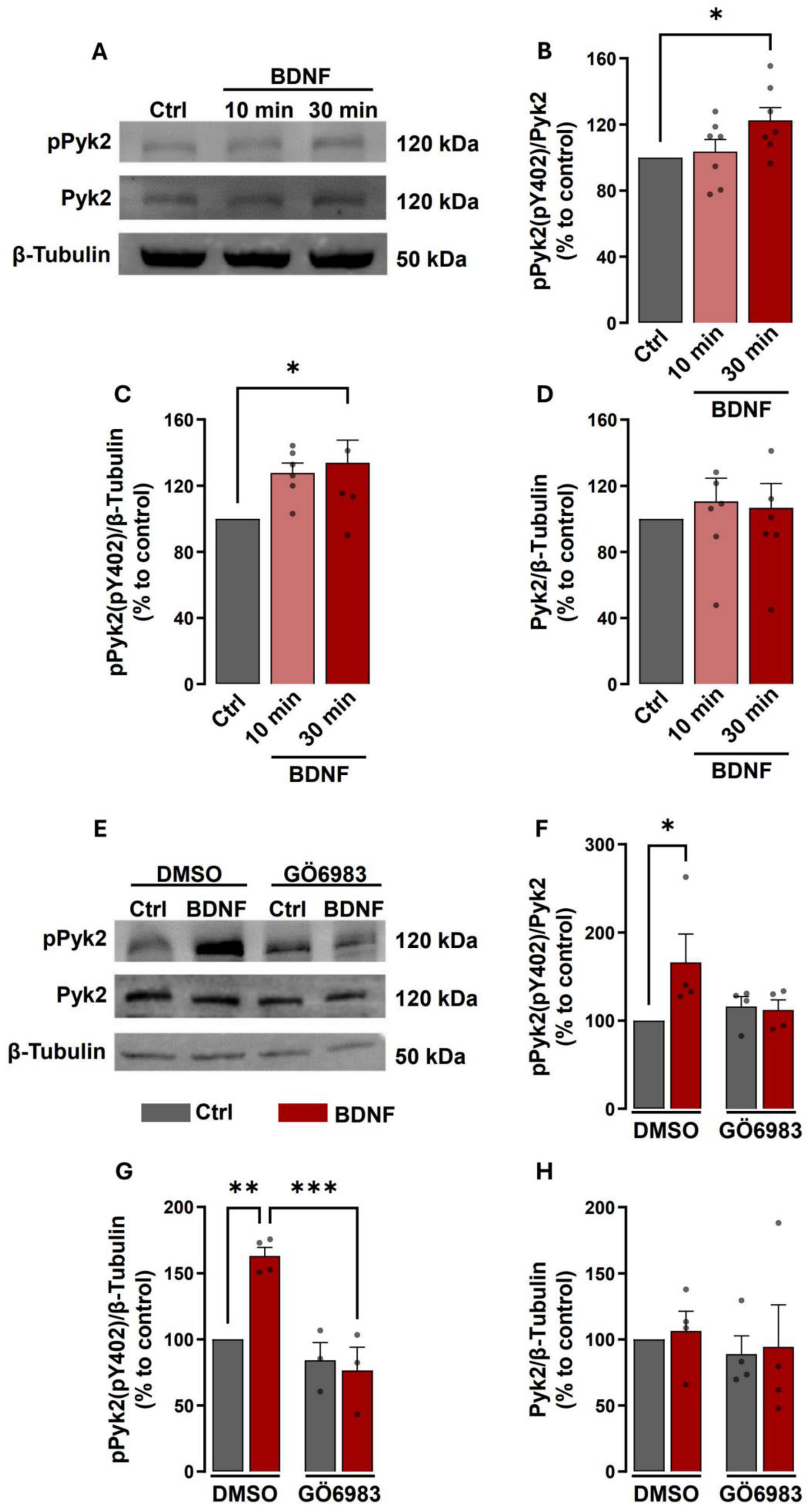
BDNF-induced activation/phosphorylation of Pyk2 is dependent on PKC activation. Representative western blots and analysis (**A**, **E**). Hippocampal neurons (14 - 15 DIV) were stimulated or not with BDNF (50 ng/mL) during 10- and 30-min (**B**, **C**, **D**) and with or without GÖ6983 where indicated (**F**, **G**, **H**). Proteins were extracted then resolved and analyzed by immunoblot, using antibodies against pPyk2, Pyk2, and β-Tubulin. In this analysis, the pPyk2 levels were normalized to β-tubulin and total Pyk2. Data are means ± SEM of at least three different experiments, performed in independent preparations. *p < 0.05, **p < 0.01, ***p < 0.001 by one-way analysis of variance (ANOVA) followed by Bonferroni post-test.

To investigate the role of PKC in BDNF-induced activation of Pyk2, we tested the effect of the kinase inhibitor GÖ 6983 (Gschwendt, Dieterich et al. 1996) on the phosphorylation of Pyk2 on Tyr402 induced by neurotrophin in cultured hippocampal neurons. The cells were preincubated with GÖ 6983 (100 nM) for 30 min before stimulation with BDNF for the same time period. Western blot analysis with an antibody specific for pPyk2(pY402) showed an increase in the phosphorylation of the kinase after BDNF stimulation and this effect was abolished by GÖ 6983 (Fig 3E, 3G). The same effect was observed when the ratio between pPyk2(pY402)/total Pyk2 was analyzed (Fig 3E, 3F). No significant changes were found in total Pyk2 protein levels under the same conditions (Fig 3E, 3H). These results show a role for PKC in the activation of Pyk2 following incubation of hippocampal neurons with BDNF.

Given the role of PKC as mediator of BDNF-induced regulation of Pyk2, we investigated whether this signaling pathway also plays a role in the neurotrophin-induced synaptic surface expression of GluN2B-containing NMDAR. In these experiments the synaptic surface expression of GluN2B-containing NMDAR was evaluated in cultured hippocampal neurons as described above, and the cells were stimulated with BDNF 30 min or maintained under control conditions. Images were analyzed for the number, intensity, and area of total and synaptic GluN2B puncta per dendritic length. Incubation of cultured hippocampal neurons with the PKC inhibitor GÖ 6983 abrogated the effects of BDNF in the upregulation of the total number of puncta of surface GluN2B staining (Fig. 4A, 4B), the area (Fig. 4A, 4C) and intensity (Fig. 4A, 4D) of those puncta. Similarly, PKC inhibition with GÖ 6983 blocked the BDNF-induced upregulation in the number of synaptic surface GluN2B puncta (Fig. 4A, 4E), as well as the area (Fig. 4A, 4F) and intensity (Fig. 4A, 4G) of those puncta. These results indicate that the regulation of synaptic expression of GluN2B-containing NMDARs upon BDNF stimulation is mediated by PKC.

**Figure 4.**
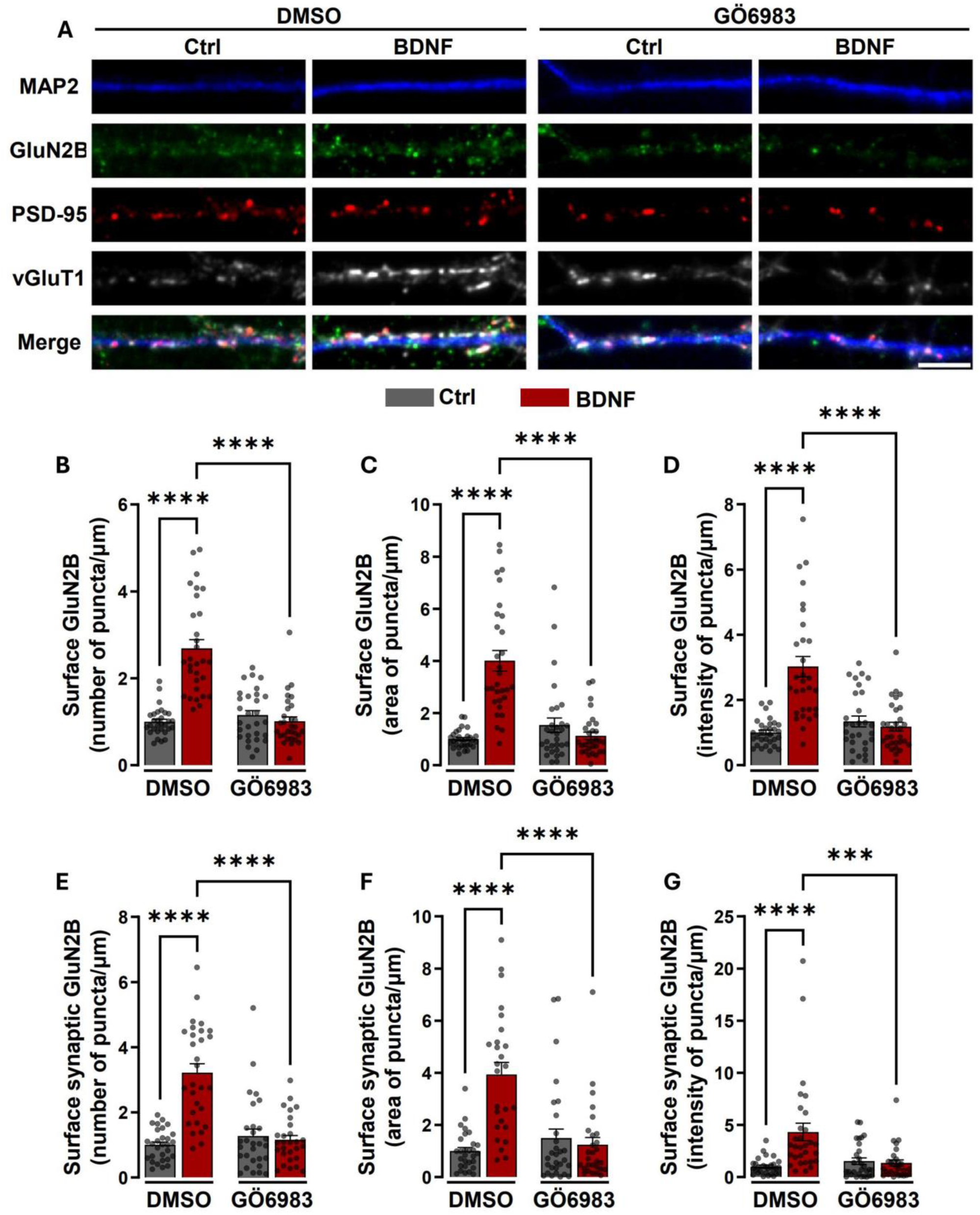
BDNF-induced increase in the synaptic expression of GluN2B-containing NMDAR is sensitive to PKC inhibition. (**A**) Representative images of hippocampal neurons (DIV 14 - 15) pre-incubated with GÖ 6983 (100 nM) or vehicle (DMSO; 1:1000 dilution for 40 min) and then either maintained under the same conditions or stimulated with BDNF (50 ng/ml for 30 min), as indicated. Neurons were live-immunoassayed for GluN2B using an antibody against an extracellular epitope in the GluN2B N-terminus, fixed, and then further immunoassayed for PSD-95, vGlut1, and MAP2. Scale bar, 5 μm. Images illustrated in (**A**) were analyzed for the total number (**B**), area (**C**), and intensity (**D**) of surface GluN2B puncta per µm. Synaptic (PSD-95- and vGlut1-colocalized) surface GluN2B number (**E**), area (**F**), and intensity (**G**) of puncta per µm of excitatory synapses (number of puncta PSD-95–vGlut1 colocalized per µm), were also analyzed. Data are normalized to the mean of the DMSO control and are the means ± SEM of 28 - 30 cells per condition, in at least three independent experiments performed in different preparations. ***p < 0.001, ****p < 0.0001 by one-way analysis of variance (ANOVA) followed by Bonferroni post-test.

### GluN2B-containing NMDAR contribute to the BDNF-induced facilitation of LTP in hippocampal CA1 synapses

We next hypothesized that the BDNF-induced synaptic accumulation of GluN2B-containing NMDAR could mediate the effects of the neurotrophin in the plasticity of glutamatergic synapses. In most excitatory synapses, activation of NMDAR induces classical forms of long-term potentiation or depression (LTP/LTD) (Bliss and Collingridge 1993, Malenka and Bear 2004). There are also several reports showing that GluN2B-containing NMDAR is involved in the consolidation of hippocampus-dependent memory (Halt, Dallapiazza et al. 2012, Sun, Cai et al. 2016, Baez, Cercato et al. 2018, Radiske, Gonzalez et al. 2021). However, the specific role of GluN2B in synaptic plasticity and memory formation is not fully understood. Moreover, several pieces of evidence point to a fundamental role for endogenous BDNF in LTP of hippocampal glutamatergic synapses (Leal, Bramham et al. 2017, Sasi, Vignoli et al. 2017, Kowianski, Lietzau et al. 2018, Rauti, Cellot et al. 2020). Furthermore, exogenous BDNF was also demonstrated to facilitate LTP (e.g. (Diogenes, Costenla et al. 2011, Santos, Mele et al. 2015)). To investigate the role of GluN2B-containing NMDAR on the facilitatory effect of BDNF upon LTP, we used a θ-burst protocol to induce LTP in the CA1 region of hippocampal slices. This protocol of stimulation has been shown to mimic the rhythm of hippocampal neurons in awake, behaving animals (Bland 1986, Larson and Munkacsy 2015). In each slice we stimulated two independent pathways, one to evaluate LTP in the absence of the drug under test and another to evaluate LTP in the presence of the drug under test (see Methods). As expected (Fontinha, Diogenes et al. 2008), the θ-burst stimulus applied in the presence of BDNF (20 ng/mL) induced a robust LTP magnitude, which was significantly higher (P < 0.01) than that obtained in the absence of BDNF. To assess the role of GluN2B in the BDNF-induced LTP facilitation, Co 101244 (1 µM), a GluN2B specific inhibitor (Higgins, Ballard et al. 2003), was added to hippocampal slices at least 20 min before the first LTP induction and remained in the bath up to the end of the experiments. In the presence of Co 101244 (1 µM), the facilitatory effect of BDNF upon LTP was totally prevented (Fig 5G-I). Incubation with Co 101244 alone did not affect the induction and maintenance of LTP, as compared with the recordings made in the same slices in another pathway before addition of the inhibitor (Fig. 5D-5F). Taken together, these results indicate that the effect of BDNF on the induction and early phases of LTP maintenance (60 min), in the CA1 region of hippocampal slices, is mediated by GluN2B-containing NMDAR.

**Figure 5.**
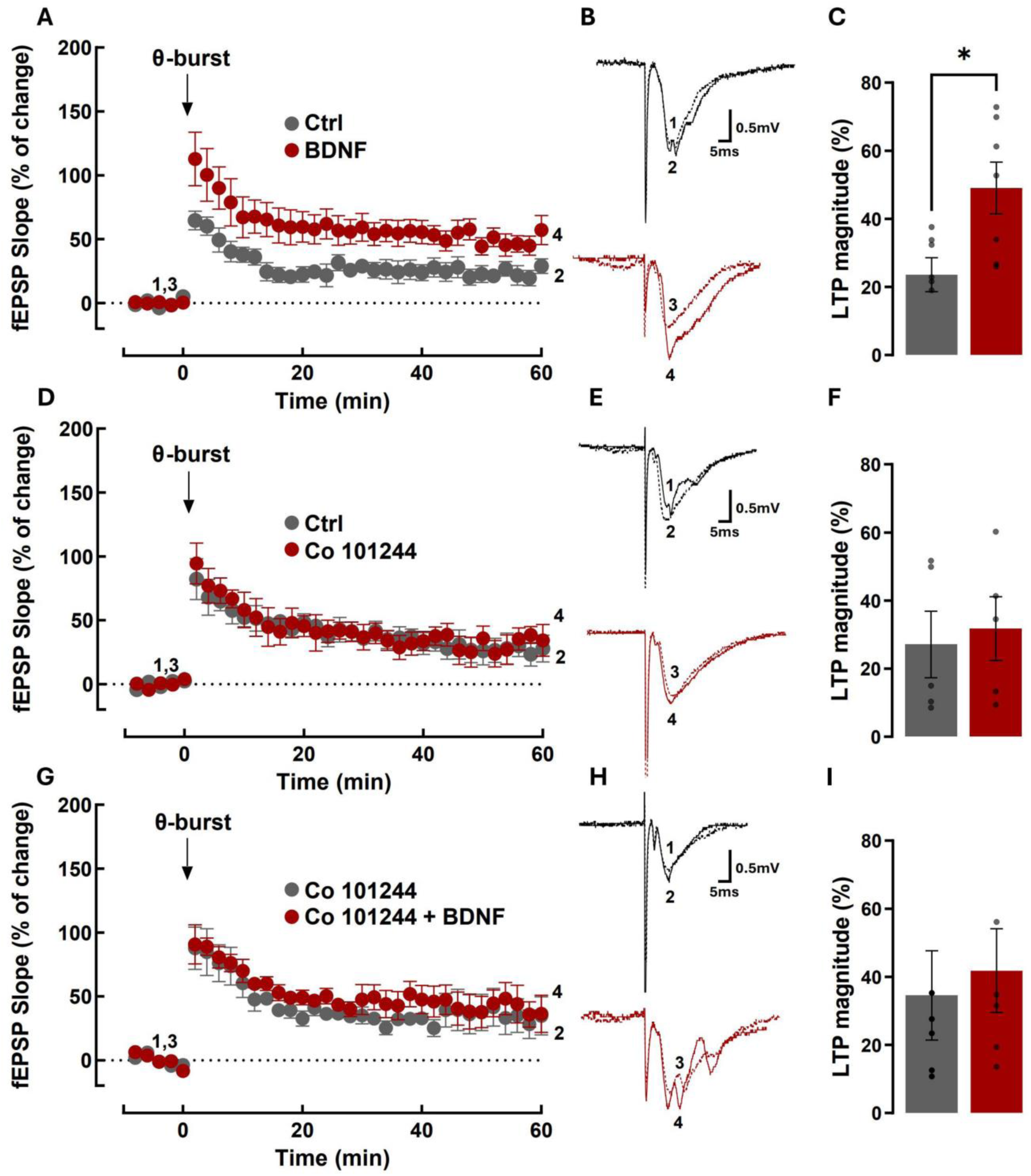
BDNF-induced enhancement of LTP is antagonized by GluN2B-containing NMDAR inhibition. (**A**) Averaged time course changes in fEPSP slope induced by a θ-burst stimulation (% of change) in the absence (black dots) or in the presence of 20 ng/ml BDNF (red dots) (**A**, **C** n= 7; **G**, **I** n= 6) or CO101244 alone (**D**, **F** n= 5). Note that in G-I BDNF was tested in the presence of CO101244. The ordinates represent normalized fEPSP slopes, where the averaged slopes recorded for 10 min before burst stimulation were set to 0%, and the abscissa represents the recording time. Representative traces (**B**, **E,** and **H**). Each trace is the average of consecutive responses obtained at the indicated periods in the time-course panel on the left, i.e. before (1 and 3) or 60 min after (2 and 4) LTP induction, and is composed of the stimulus artifact, followed by the presynaptic volley and the fEPSP. Traces 1 and 2 were obtained in the absence of BDNF and traces 3 and 4 in its presence from a second pathway in the same slice. Traces recorded from the same pathway before and after LTP induction are superimposed. (**C**, **F**, **I**) Bar graphs show the magnitude of LTP (change in the fEPSP slope over time) induced by θ-burst stimulation in relation to pre-burst values (0%) in the same hippocampal slices. Statistical analysis for the LTP magnitude was performed by the *t*-test. (*p < 0.05).

### TrkB signaling is required for the upregulation of synaptic GluN2B expression in status epilepticus

To determine whether BDNF signaling is also coupled to the regulation of the synaptic expression of GluN2B in status epilepticus we analyzed the synaptic surface abundance of this NMDAR subunit in rats subjected to the pilocarpine model of temporal lobe epilepsy (TLE), under control conditions and in animals treated with ANA-12, a TrkB receptor inhibitor (Fig. 6A), thus allowing to assess the role of endogenous BDNF. The surface expression of GluN2B was assessed as described in the experiments illustrated in Fig. 1, and representative images are shown in Fig. 6B. The integrated density of GluN2B staining was increased in animals treated with pilocarpine, and this effect was inhibited by administration of ANA-12 (Fig. 6B, C). In contrast, TrkB inhibition was without effect on the alterations in the pattern of GluN2B staining regarding the mean gray value of GluN2B immunoreactivity that colocalized with the synaptic markers (Fig. 6B, D). In addition, administration of ANA-12 under control conditions did not change GluN2B staining in hippocampal synaptoneurosomes.

**Figure 6.**
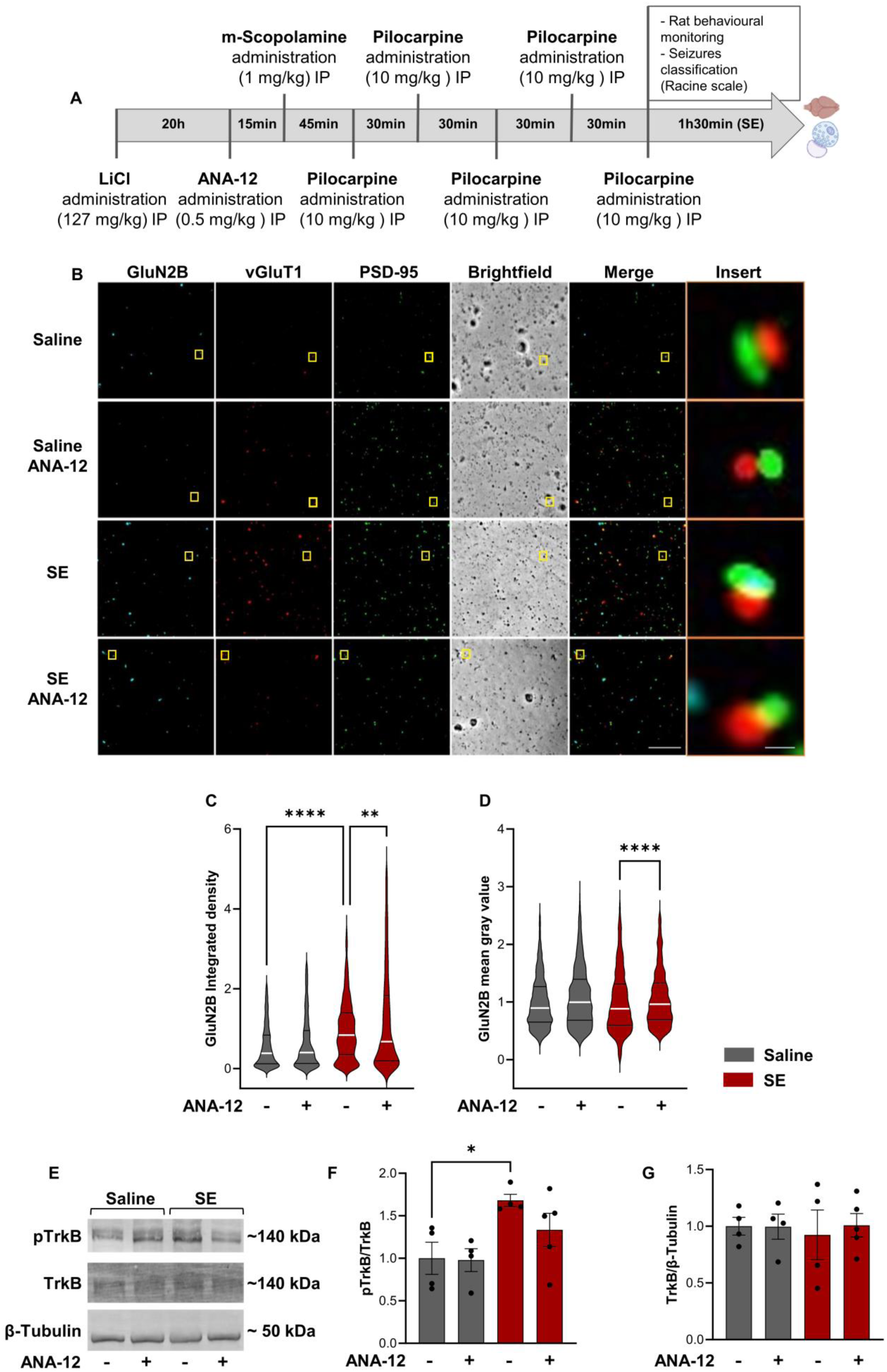
The increase in the expression of GluN2B-containing NMDAR in hippocampal synaptoneurosomes during epileptogenesis requires TrkB signaling. (**A**) Experimental design for the lithium-pilocarpine model of Status Epilepticus. (**B**) Representative images of hippocampal synaptoneurosomes (prepared from adult Sprague-Dawley rats 6-8 weeks old, treated with saline, saline and ANA-12, Pilocarpine or Pilocarpine and ANA-12). Synaptoneurosomes were live-immunoassayed for GluN2B using an antibody against an extracellular epitope in the GluN2B N-terminus and immunoassayed for vGlut1 and PSD-95. Merge scale bar, 10 μm. Insert scale bar, 0.5 µm. Images illustrated in (**B**) were analyzed for the GluN2B integrated density (**C**) and GluN2B mean gray value (**D**). Data are the means ± SEM of 779 - 840 synaptoneurosomes per condition, from at least four animals for each experimental condition. ****p < 0.0001, **p<0.01 as determined by Kruskal Wallis’s test and Dunn’s multiple comparisons test. Representative western blot and analysis (**E-G**). Proteins were extracted from synaptoneurosomes prepared from the same animals used in the immunocytochemistry experiments. For the immunoblot, antibodies against pTrkB, total TrKB and β-Tubulin were used. In this analysis, pTrkB levels were normalized to total TrkB levels and total TrkB levels were normalized to β-tubulin. Data are means ± SEM of at least four animals for each experimental condition. *p < 0.05, by one-way analysis of variance (ANOVA) followed by Dunnett’s post-test.

In another set of experiments we assessed TrkB receptor activity in hippocampal synaptoneurosomes under the same experimental conditions, by measuring the phosphorylation of the receptor on Tyr816. The total levels of TrkB were also analyzed by western blot, and the results were expressed as a ratio of pTrkB/TrkB (Fig. 6E, F). An upregulation of the ratio pTrkB/TrkB was observed in hippocampal synaptoneurosomes from rats subjected to pilocarpine treatment when compared with animals treated with vehicle. These results confirm the activation of TrkB receptors in hippocampal synaptosomes under the conditions described above, which showed TrkB mediated alterations in the synaptic expression of GluN2B-containing NMDAR. Accordingly, no significant alteration in the pTrkB/TrkB ratio was observed in the animals subjected to pilocarpine treatment after administration of the TrkB receptor inhibitor, ANA-12. Furthermore, ANA-12 had no effect on the pTrkB/TrkB ratio when administered to control animals treated with saline only (Fig. 6E, F). In addition, no alterations in total TrkB protein levels were observed in hippocampal synaptoneurosomes under the experimental conditions used (Fig. 6E, G).

## Discussion

Plasticity of glutamatergic synapses in the hippocampus, including LTP induced by theta-burst presynaptic stimulation and the alterations associated with epileptogenesis, has been associated with an upregulation in the synaptic expression of GluN2B-containing NMDAR, but the molecular mechanisms underlying these changes have not been elucidated. The results obtained in this work point to a key role for BDNF in the synaptic expression of GluN2B in hippocampal synapses under conditions characterized by an activity-dependent upregulation of glutamatergic activity. This hypothesis is supported by several findings: (i) BDNF induced the surface expression of GluN2B-containing NMDAR in hippocampal synaptoneurosomes; (ii) incubation of cultured hippocampal neurons with BDNF increased the synaptic surface accumulation of GluN2B by a mechanism mediated by PKC; (iii) BDNF facilitated LTP induced by θ-burst stimulation of hippocampal CA1 synapses through a mechanism that requires GluN2B activity; (iv) inhibition of TrkB decreased the early synaptic surface accumulation of GluN2B in hippocampal synaptoneurosomes isolated from the hippocampus of rats subjected to the pilocarpine model of temporal lobe epilepsy. The GluN2B-containing NMDAR are characterized by an enhanced Ca^2+^-permeability and charge transfer, as well as by a slower rate of deactivation, rise and decay times when compared with the receptors with GluN2A subunits (Vieira, Yong et al. 2020). Therefore, these changes in the molecular composition of NMDAR at the synapse are expected to have a significant impact in the activation of downstream signaling.

BDNF induced a time-dependent synaptic surface accumulation of GluN2B-containing NMDAR in hippocampal synaptoneurosomes. The effects of BDNF in a synaptic fraction isolated from adult rats resemble the effects of the neurotrophin in the synaptic accumulation of GluN2B in cultured hippocampal neurons, showing that synaptoneurosomes are a valuable model to study the regulation of the synaptic expression of NMDAR. Biotinylation studies have revealed that BDNF promotes the surface expression of GluN2A and GluN2B in cultured hippocampal neurons, albeit with distinct kinetics (Caldeira, Melo et al. 2007). In addition, BDNF upregulates the synaptic expression and function of GluN2B-containg NMDAR in cultured hippocampal neurons by a mechanism dependent on the local synthesis of the cytosolic tyrosine kinase Pyk2 using transcripts transported along dendrites by the RNA binding protein heterogenous nuclear ribonucloprotein K (hnRNP K) (Leal, Comprido et al. 2017, Afonso, De Luca et al. 2019). In this work we identified an additional player in the mechanism underlying the regulation of GluN2B synaptic expression by BDNF. Thus, inhibition of PKC abrogated the effect of BDNF in the synaptic surface accumulation of GluN2B and this effect was correlated with an impairment in the BDNF-evoked phosphorylation of Pyk2 on Tyr402 which is important in the activation of downstream signalling (Das, Pal et al. 2021, de Pins, Mendes et al. 2021). Since TrkB receptors are coupled to the activation of phospholipase Cγ with a consequent activation of PKC (Leal, Comprido et al. 2014), the downstream regulation of Pyk2 is likely to account for the effects of BDNF on the surface expression of GluN2B-containing NMDAR. Indeed, dimerization and activation of Pyk2 is followed by stimulation of the Src family kinase Fyn (Dikic, Tokiwa et al. 1996, Walkiewicz, Girault et al. 2015) which phosphorylates GluN2B (Nakazawa, Komai et al. 2001, Yang, Trepanier et al. 2012). PKC activation has also been shown to be an upstream regulator of Pyk2 following NMDAR stimulation which is essential for the neuronal synaptic transmission (Sun, Zhang et al. 2016, de Pins, Mendes et al. 2021, Mastrolia, Al Massadi et al. 2021). Such mechanism may contribute to amplify the signaling cascade to upregulate the activity of glutamatergic synapses.

Inhibition of GluN2B-containing NMDAR with Co 101244 abrogated the facilitatory effects of BDNF on LTP of hippocampal CA1 synapses without affecting synaptic potentiation in the absence of exogenous neurotrophin. These results show that BDNF acts specifically by functional recruitment of a population of NMDAR containing GluN2B subunits which do not contribute to LTP with the θ-burst protocol used here. Protein kinase C signaling may mediate the synaptic recruitment of GluN2B-containing NMDAR to mediate the effect of BDNF on the facilitation of LTP of hippocampal synapses, as suggested by the evidence obtained in the experiments performed in cultured hippocampal neurons. Accordingly, hippocampal LTP is supported by presynaptic and postsynaptic tyrosine receptor kinase B-mediated phospholipase Cγ signalling (Gartner, Polnau et al. 2006). We also found that PKC mediates the phosphorylation (activation) of Pyk2 in cultured hippocampal neurons stimulated with BDNF. This signalling pathway may mediate the effects of BDNF in LTP since Pyk2-deficient mice exhibited an impairment in LTP in CA1 synapses as well as a cognitive impairment in hippocampal-related tasks (Giralt, Brito et al. 2017).

Hippocampal synapses in adult rats are enriched in GluN2A-containing NMDAR (Barria and Malinow 2002). Therefore, BDNF-TrkB signaling likely induces a rapid change in the molecular composition of these receptors by increasing the abundance of GluN2B subunit, with a consequent alteration in the receptor electrophysiological and signaling properties (Yashiro and Philpot 2008). In particular, the localization of GluN2A and GluN2B containing NMDAR in distinct nanodomains in hippocampal synapses (Kellermayer, Ferreira et al. 2018) may allow their coupling to the activation of distinct intracellular effector mechanisms. The BDNF-induced facilitation of LTP was previously shown to be mediated by a decrease of the proteasome activity (Santos, Mele et al. 2015). Since Ca^2+^-entry through NMDAR is coupled to the inhibition of the proteasome activity (Caldeira, Curcio et al. 2013), this mechanism may account for the effects of BDNF in the facilitation of LTP reported here. Inhibitors of protein synthesis also blocked BDNF-induced facilitation of CA1 synapses (Kang and Schuman 1996) and accordingly *in vivo* studies showed that protein synthesis is also required for induction and consolidation of long-term potentiation (LTP) elicited by local infusion of BDNF in the dentate gyrus of anesthetized rats (Messaoudi, Kanhema et al. 2007). The simultaneous effects of BDNF in the regulation of translation and protein degradation by the proteasome is expected to provide a tight control of the synaptic proteome in those forms of synaptic potentiation that are mediated by the neurotrophin.

An upregulation in the surface expression of GluN2B was observed in hippocampal synaptoneurosomes from rats subjected to the lithium-pilocarpine model of temporal lobe epilepsy, at the stage of status epilepticus. A previous study using field excitatory postsynaptic potential recordings also reported an increase in functional responses mediated by GluN2B-containing NMDAR in the lateral perforant path synapses with hippocampal dentate granule neurons at the chronic phase of epilepsy in the pilocarpine model (Klatte, Kirschstein et al. 2013). However, functional studies do not allow distinguishing the synaptic alterations from an upregulation in the surface expression of the receptors and those mediated by posttranslational modifications. Importantly, in this study we found that the increased synaptic surface expression of GluN2B in hippocampal synaptoneurosomes at the stage of status epilepticus in the lithium-pilocarpine model of temporal lobe epilepsy is mediated by TrkB receptors. This is in accordance with the results showing a rapid upregulation in BDNF under the same conditions (Tongiorgi, Armellin et al. 2004), and with the effect of a peptide that uncouples TrkB from PLCγ1 which prevents epilepsy induced by status epilepticus after kainate infusion in the amygdala (Gu, Huang et al. 2015). Administration of the peptide following a seizure also reverted a subset of animals to an earlier state of epileptogenesis in a modified kindling model in which seizures were induced through amygdala stimulation (Krishnamurthy, Huang et al. 2019). The increased Ca^2+^ permeability resulting from the synaptic expression of GluN2B-containing NMDAR (Yashiro and Philpot 2008) may contribute to further enhance the exocytotic release of BDNF (Brigadski and Lessmann 2020) to further potentiate the glutamatergic synapses. In addition, the entry of Ca^2+^ through GluN2B-containing NMDAR may also lead to the activation of downstream signaling coupled to the plasticity of spines (Jean, Carton et al. 2023).

In conclusion, in this work we have shown that BDNF signaling is coupled to the upregulation of glutamatergic synapses by increasing the activity of GluN2B-containing NMDAR. The latter responses account for the effects of BDNF in the facilitation of glutamatergic synapses in the CA1 region of the hippocampus. An upregulation in the synaptic expression of GluN2B in hippocampal synapses was also observed in the lithium-pilocarpine model of temporal lobe epilepsy. These effects of BDNF in the potentiation of glutamatergic synapses are expected to further enhance the release of the neurotrophin, and may therefore act as a positive feedback loop. Such mechanisms are expected to remain within physiological levels in plasticity-related phenomena, but when exacerbated may contribute to epileptogenesis.

